# Lung adenocarcinoma cell-derived exosomal miR-328 promote osteoclastogenesis by targeting Nrp2

**DOI:** 10.1101/2021.07.08.451611

**Authors:** Chengcheng Zhang, Jingru Qin, Lu Yang, Zhiyao Zhu, Jia Yang, Wan Su, Haibin Deng, Zhongqi Wang

## Abstract

Bone metastasis of lung cancer and detailed mechanisms are still elusive, and the roles of exosomes derived from lung adenocarcinoma cells in this process have attracted much attention. In this study, we found that A549 cell-derived exosomes (A549-Exos) promoted osteogenesis and bone resorption in vitro. Furthermore, A549-Exos target bone in vivo and promoted bone resorption in vivo. Mechanistically, A549-Exosomal miR-328 promoted bone resorption by targeting Nrp2 and A549-Exos^miR-328 Inhibitors^ inhibited bone resorption *in vivo*. Thus, A549-Exosomal miR-328 promote osteoclastogenesis by targeting Nrp2 and A549-Exos^miR-328 Inhibitors^ may serve as a potential nanomedicine for the treatment of bone metastasis.

## Background

Bone metastasis is the most frequent site of lung cancer metastasis^1^ and most lung cancer bone metastasis can be classified into osteolytic bone metastasis^2^, which can cause pain, pathological, spinal instability, spinal cord compression and hypercalcemia and result in a decline in the quality of life^3,4^. However, lung cancer cells can not directly lead to osteolytic metastasis by proteolytic enzymes. Instead, lung cancer cells derived circulating miRNAs and exosomes played important roles by activating osteoclasts^5,6^. Thus, we focus on the impact of lung cancer derived exosomes on osteoclast differentiation.

Exosomes are small cup or sphere-shaped vesicles, and their diameters primarily ranged from 80 nm to 150 nm, that contains mRNAs, miRNAs^7^, lipids^8^ and proteins^9^. It has been reported that the role of exosomes in cellular-communication by the transfer of proteins^10^, bioactive lipids and miRNAs^11^. Lung cancer derived exosomes have been attracted attention for its role in bone metastasis and prognosis^12^. Furthermore, exosomes may be potentially used as biomarkers, nanoparticle drugs and carriers of antitumor drugs. Recently, numerous investigations indicated that exosomal miRNAs played important roles in tumor growth, invasion and metastasis^13^. Therefore, finding the key exosomal miRNAs in the bone metastasis could be beneficial for treatment of bone metastasis and exosomal miRNAs may serve as a potential therapeutic target for bone metastasis^14^.

In this study, we found that A549 cells derived exosomes are important extracellular vesicles in osteoclast differentiation and in bone metastasis. Furthermore, we found that A549-Exos promoted bone resorption in vitro and in vivo. We furthermore explored its mechanism and found that A549-Exosomal miR-328 promoted bone resorption by targeting Nrp2. Therefore, we generated A549-Exos loaded in miR-328 inhibitors and found that bone resorption was prevented by injecting A549-Exos^miR-328 Inhibitors^. Therefore, we focused on the roles of A549 cells derived exosomes in bone resorption in this study.

## Methods

### Cell culture

A549 cell line was purchased from the American Type Culture Collection (ATCC) and cultured in Dulbecco’s modified Eagle’s medium (DMEM) (Gibco, USA), supplemented with 10% fetal bovine serum (FBS) (Gibco, USA). The Raw264.7 cell line was cultured in DMEM (Gibco) supplemented with 10% FBS. We cultured cells at 37 °C in a 5% CO_2_ humidified incubator.

### Exosome isolation and characterization

A549 cells were cultured in fresh complete medium containing EVs-free FBS (Thermo, USA) for 48 h. Then, we collected the conditioned medium and centrifuged them at 1500 g for 15 minutes at 4°C. After filtering through a 0.22 mm filter (Millipore, Billerica, USA). The supernatant was ultracentrifuged twice at 110 000 g for 1 hour at 4°C. Then, we resuspended the pellets in a right amount of PBS and stored A549-Exos at –80°C for later use. We observed the morphology of A549-Exos by transmission electron microscope (TEM). The size distribution of A549-Exos was determined by using Nanosizer™ instrument (Malvern Instruments, Malvern, UK). The expression of exosomal surface markers (CD9, TSG101) was analyzed by western blot (WB).

### Western blot (WB) analysis

Cells or A549-Exos were lysed in a RIPA lysis buffer supplemented with protease inhibitor cocktail (CST). Then, we quantified total protein concentration by using the BCA protein assay kit (Thermo, USA). Western blotting was performed as previously described^15^. Primary antibodies were used in this study as follows: CD9 (CST, USA), HSP70 (CST, USA), GAPDH (CST, USA) and OPG (CST, USA). All the secondary antibodies (1:5000) were acquired from Abcam.

### In vitro Osteoclastogenesis assays of RAW264.7 cells

We seeded RAW264.7 cells on 48-well plates ((1× 10^4^)) in D-MEM (Gibco, USA) supplemented with 10% FBS (Gibco, USA). M-CSF (30ng/ml) and RANKL (100ng/ml) were used to promote osteoclast maturation for 7 days. Then, we performed TARP-staining according to general protocols. We considered TRAP positive cells (more than 5 nucleus) considered as mature osteoclasts.

### Luciferase reporter assay

We constructed the Nrp2 3′-UTR luciferase reporter construct by amplifying the mouse Nrp2 mRNA 3′-UTR sequence (including the predicted miR-328 binding site) by PCR. Then, the PCR products were purified and cloned into the XbaI site of the pGL3-promoter vector (Promega) immediately downstream of the stop codon of luciferase. The Nrp2 mutations were prepared by using a Quick-Change Site-Directed Mutagenesis Kit (Stratagene) to get MUT-pGL3-Nrp2, which is confirmed by sequencing. RAW264.7 cells were transfected with either WT or mutant pGL3 construct and 40 ng of pRL-TKRenilla-luciferase plasmid and miR-328 mimics or inhibitors for 48 h. Luciferase activities were measured as previously described^15^.

### Real-time PCR (RT-PCR)

Total RNA of RAW264.7 cells or A549-Exos were isolated by TRIzol reagent (Life Technologies, USA) and synthesized complementary DNAs (cDNAs). Then, Quantitative reverse transcriptase PCR (qRT-PCR) was performed as previously described^15^. The primer sequences used for qRT-PCR were as follows: miR-328: forward, 5’-CGG GCC TGG CCC TCT CTG CC-3’; reverse, 5’-CAG CCA CAA AAG AGC ACA AT-3’; Ctsk: forward, 5’-CTC GGC GTT TAA TTT GGG AGA-3’; reverse, 5’-TCG AGA GGG AGG TAT TCT GAG T-3’; TRAF-6: forward, 5’-AAG GTG GTG GCG TTA TAC TGC-3’; reverse, 5’-CTG GCA CAG CGG ATG TGA G-3’ and GAPDH: forward, 5’-AAT GGA TTT GGA CGC ATT GGT-3’; reverse, 5’-TTT GCA CTG GTA CGT GTT GAT-3’;

### Animal studies

All animal experiments were conducted in accordance with the National Institute of Health (NIH) Guide for the Care and Use of Laboratory Animals, with the approval of Longhua hospital (No. LHERAW-19038). C57BL/6 mice (6-8 weeks old) were obtained from the Animal Center of Longhua hospital. The mice were housed in barrier housing conditions in the Animal Center of Longhua hospital. For delivery of A549-Exos or A549-Exos^miR-328 Inhibitors^, 100ng/ml in 1 ml PBS was injected into tail vein once every 3 days for 4 weeks. The mice were sacrificed at 4 weeks after the treatment.

### Micro-CT analysis

femur samples dissected from mice were scanned and analyzed through micro-CT (Quantum GX, PE, USA). We performed the micro-CT scans under the same conditions: voltage 100 kV, current 80 μA, spatial resolution 12 μm, scanning 500 continuous sections. The data was collected and analyzed automatically through computer software (Skyscan) to analyze the number of trabecular bones (Tb.N), trabecular bone thickness (Tb.Th), trabecular bone space (Tb.Sp), bone volume fraction (BV/TV) and three-dimensional reconstruction.

### Exosomes labeling and tracking in vivo

All animal experiments were conducted in accordance with the National Institute of Health (NIH) Guide for the Care and Use of Laboratory Animals, with the approval of Longhua hospital (No. LHERAW-19038). To visualize the target organ and the metabolism process of exosomes, we labelled A549-Exos according to the instructions of the manufacturer (Thermo, USA). First, we incubated 100ug of exosomes with Vybrant DID (1:1000 in PBS) in the dark for 30 mins, and labelled exosomes were washed in PBS, with centrifugation at 100,000 × g for 60 min. Then, the labelled exosomes were injected into the tail vein of mice (8 weeks old, n=3). The mice were harvested at 3, 6, 12 hours after injection. The mean intensities for DID signals were quantified from four random visual fields for each section.

### HE & TRAP staining

For histological staining, we dissected femora from different groups and fixed them in 4% paraformaldehyde for 24 h, then decalcified them in ethylene diamine tetraacetic acid (EDTA; pH=7.4) for 28 days, dehydrated them through ethanol and embedded in paraffin. Then, we cut them into 10 μm thick longitudinally oriented sections and performed hematoxylin and eosin (H&E) staining and TRAP staining as described previously^15^.

### Statistical analysis

All data are shown as means ± standard deviation (SD) from at least three independent experiments. Comparisons were performed using t test and one-way ANOVA as appropriate. P < 0.05 was considered statistically significant.

## Results

### Characteristics of the isolated A549-Exos

To investigate the role of lung adenocarcinoma cell in osteoclastogenesis, we first determined whether lung adenocarcinoma cell could affect the differentiation of osteoclasts, RAW264.7 cells were co-cultured with A549 cells in a transwell system. The differentiation of osteoclasts was enhanced when co-culturing with A549 cells (Figure 1A). Then, we performed TEM, Dynamic light scattering analysis and Western Blot analysis to charactize the exosomes (Exos) derived from A549. A549-Exos presented a cup or sphere-shaped morphology and their diameters primarily ranged from 80 nm to 150 nm (Figure 1B, 1C). Western Blot analysis showed that A549-Exos expressed exosomal marker proteins including CD9 and TSG101 (Figure 1D). Thus, these data suggested that Osteoclastic differentiation was enhanced when co-culturing with A549 cells and A549-Exos showed characteristic features of exosomes.

**Figure 1.**
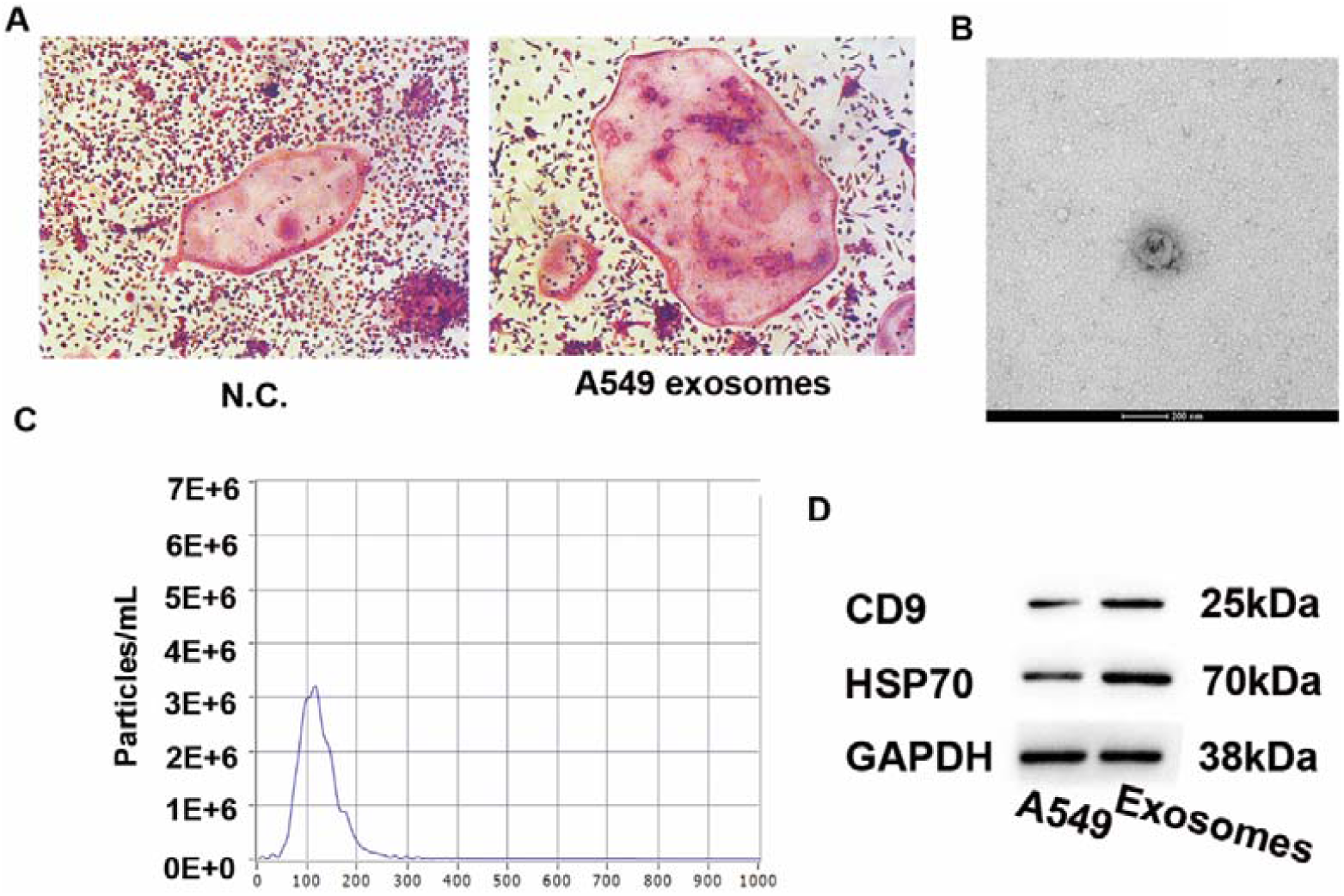
Characteristics of the isolated A549-Exos. (A) Representative images of TRAP staining osteoclasts. Scale bar: 100 μm. (B) Representative TEM image of A549-Exos (scale bar = 200 nm). (C) Representative NTA results showing the size distribution of A549-Exos. (C) Detection of A549-Exos surface markers (CD9 and HSP70) by Western Blot analysis.

### A549-Exos promoted osteoclast differentiation and bone resorption in vitro

To investigate the role of A549-Exos in osteoclastogenesis in vitro. A549-Exos were labeled with lipophilic dye Dio and assessed whether A549-Exos could be transported to Raw264.7 cells. The fluorescence microscope revealed that A549-Exos could be successfully absorbed by Raw264.7 cells (Figure 2A). Then, we treated Raw264.7 cells with A549-Exos or an equal volume of Phosphate Buffer Saline (PBS). osteoclastic differentiation was upregulated as the concentration of exosomes derived from A549 cells increased (Figure 2B, 2C). Furthermore, we found that mRNA expression of Ctsk and Traf-6 were upregulated by A549-Exos compared to its’ negative control at day 7 (Figure 2D, 2E). Taken together, these data suggested that A549-Exos promoted osteoclast differentiation in vitro.

**Figure 2.**
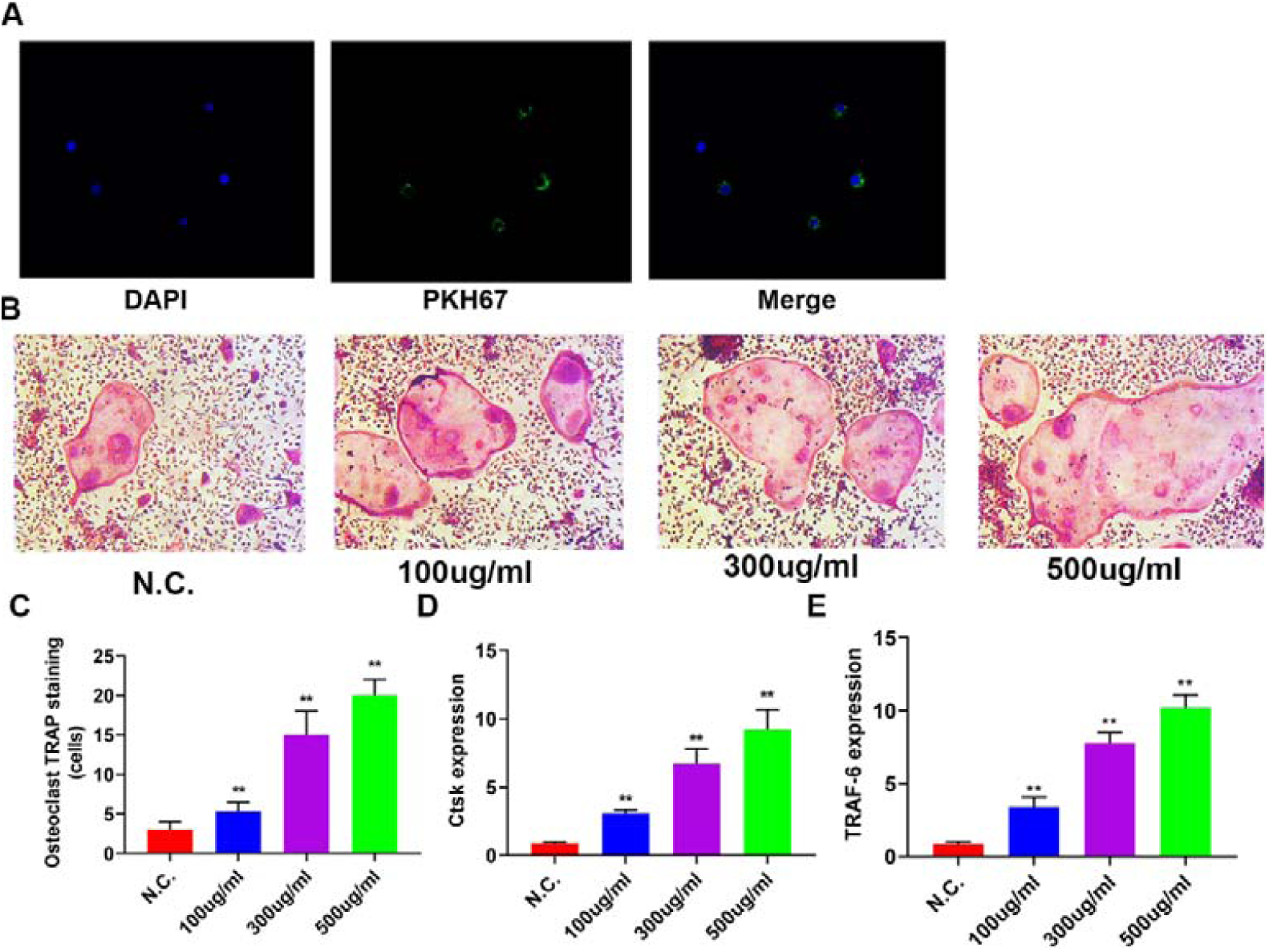
A549-Exos promoted osteoclast differentiation and bone resorption in vitro. (A) Fluorescence microscopy analysis of RAW264.7 cells treated with DiO-labeled A549-Exos for 2h. (B) Representative images of TRAP staining of mature osteoclasts treated with different concertrations of A549-Exos. Scale bar: 100 μm. (C) Quantitative analysis of positive TRAP staining osteoclasts in Figure 2B (**P<0.01). (D) Relative levels of Ctsk (left) and TRAF-6 (right) mRNA expression in Figure 2B (**P<0.01).

### A549-Exos target bone in vivo

To determine the bio-distribution of A549-Exos in vivo. The tissue distribution of DID-labeled A549-Exos (A549-Exos-DID) was examined by fluorescence molecular tomography (FMT) imaging system. A549-Exos-DID was injected into the tail vein of 8-week-old male mice for 3 h, 6 h or 12 h. 3 hours after intravenous injection, the major fluorescence signals were detected in the liver and lungs and the signals can be detected in the limbs. After 6 h, the major fluorescence signals were still accumulated in the liver and lung. Significantly, fluorescence signals in the limbs were enhanced, but still weaker than those in the liver and lung. After 12 hours, fluorescence signals still can be detected in the limbs, and fluorescence signals faded from liver and lungs. The fluorescence signals can be rarely detected in the kidney (Figure 3A). Furthermore, HE staining suggested that there was no obvious significant histologic difference in heart, liver, lung, and kidney in different groups, suggesting that A549-Exos are well-tolerated, with little to no acute, systemic toxicity (Figure S1). Taken together, A549-Exos could be targeted for bone tissues in vivo without toxicity.

**Figure 3.**
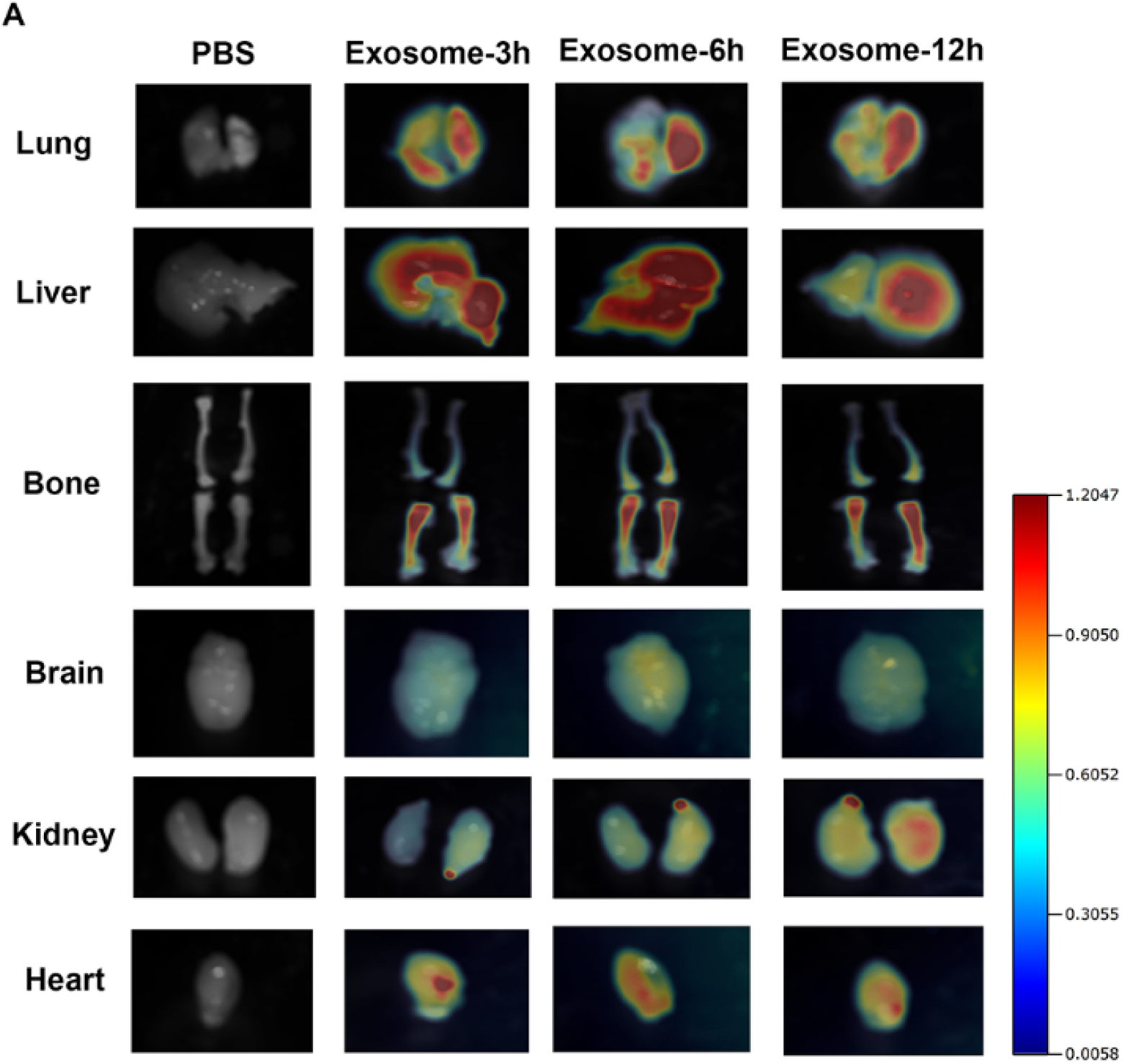
A549-Exos target bone in vivo without toxicity. (A) Representative FMT images of the near-infrared fluorescence signals in organs isolated from mice administered with PBS, A549-Exos for different periods of time (3 hours, 6 hours, and 12 hours).

### A549-Exos inhibited bone formation in vivo

To investigate the role of A549-Exos in vivo, we injected A549-Exos or an equal volume of PBS into tail vein of 8-week-old male mice third per week for 12 weeks. Microcomputed tomography (micro-CT) showed significantly lower trabecular bone volume, number, thickness, and higher trabecular separation in those mice injected with A549-Exos compared to their controls (Figure 4A, 4B-4E). Furthermore, TRAP staining showed that a greater number of osteoclasts on the trabecular bone surface in A549-Exos groups than N.C., suggesting an agonist role of A549-Exos in the osteoclast differentiation, which was consistent with our in vitro study (Figure 4F, 4H). Our results indicated that A549-Exos promoted bone resorption in vivo.

**Figure 4.**
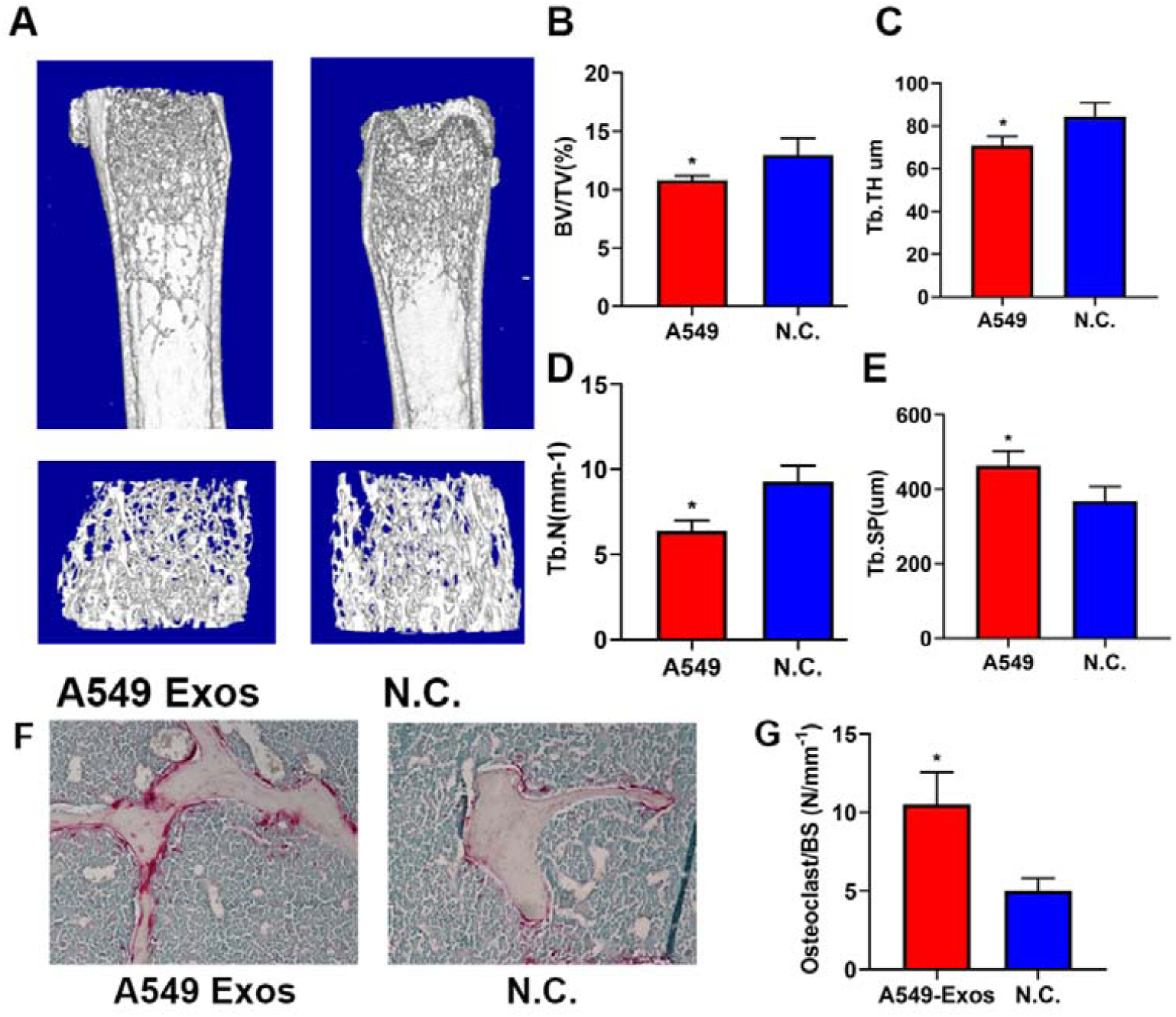
A549-Exos inhibited bone formation in vivo. (A) Representative micro-CT images in femora from A549-Exos mices and negative control group (n = 8 per group). Scale bar: 100 μm. (B, C, D, E) Quantitative analysis of trabecular microarchitecture from A549-Exos group and negative control group. (F) Representative TRAP staining of femora from A549-Exos mice and negative control group. (G) Quantitative analysis of osteoclasts per bone surface in different groups. BV/TV, trabecular bone volume per tissue volume; Tb. Th, trabecular thickness; Tb. Sp, trabecular separation; Tb. N, trabecular number. (n=8 per group) (*P<0.05).

### A549-Exosomal miR-328 inhibited bone formation by targeting Nrp2

Since exosomes may concentrate large numbers of miRNAs, which can regulate gene expression by binding 3′-untranslated regions. MiR-328 has been reported to promote osteoclast differentiation^16^. Subsequently, we examined the expression levels of miR□328 in the exosomes, and qRT-PCR showed that A549 cells derived exosomes had significantly higher expression of miR-328 in comparison with TC-1 cells derived exosomes (Figure 5A).

**Figure 5.**
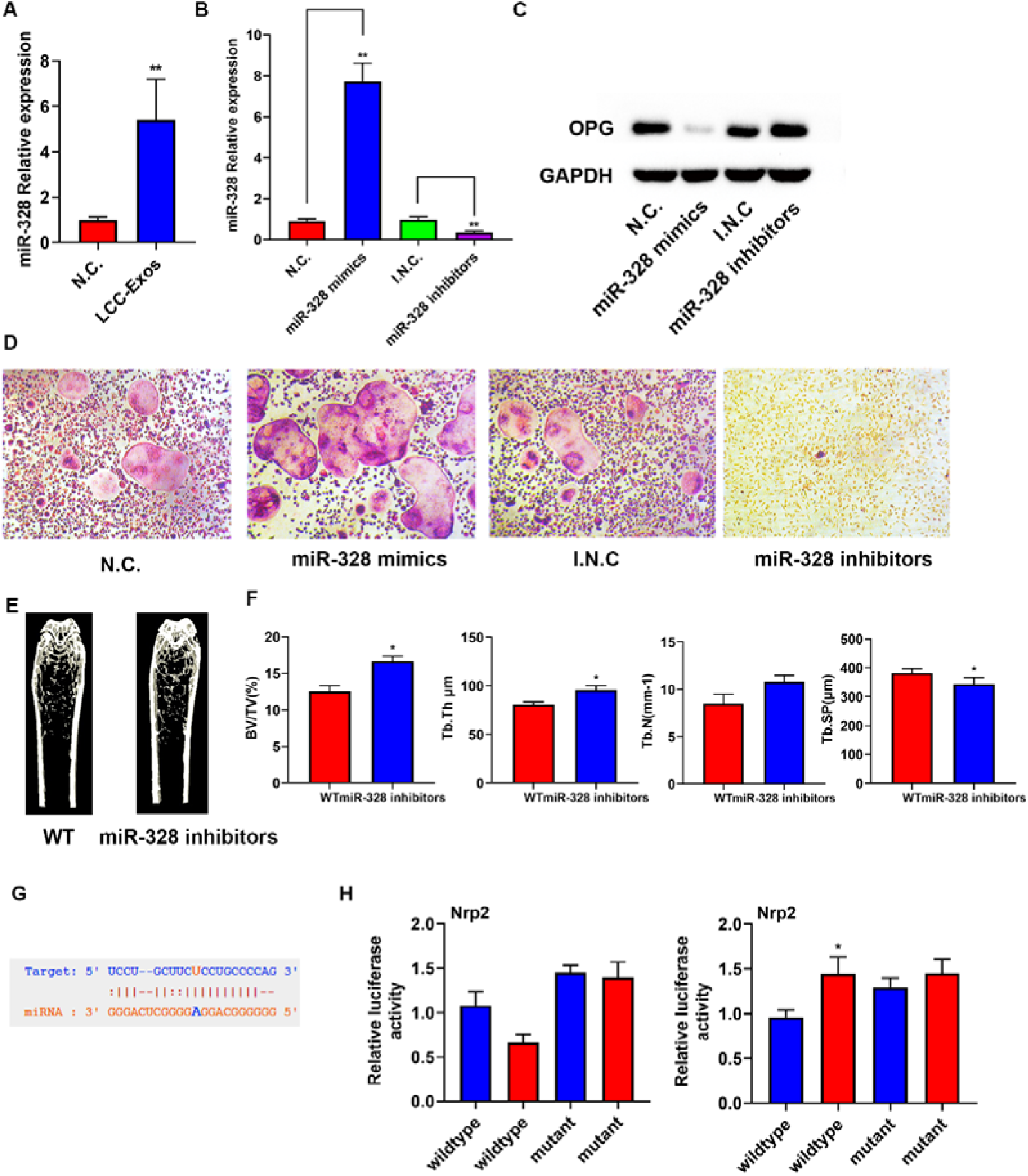
A549-Exosomal miR-328 promoted bone resorption by targeting Nrp2. (A) miR-328 relative expressions in N.C. and LCC-Exos. (B) miR-328 levels in N.C., miR-328 mimics, I.N.C. and miR-328 inhibitors were measured by qRT-PCR. (C) Western blot (WB) analysis of OPG and GAPDH in RAW264.7 cells transfected with miR-328 mimics and miR-328 or their respective negative controls. (D) Representative images of TRAP staining of mature osteoclasts treated with N.C., miR-328 mimics, I.N.C. and miR-328 inhibitors. (E) Representative micro-CT images in femora from different groups. (F) Quantitative analysis of trabecular microarchitecture from different groups. (G) Nrp2 might be the target of miR-328. (H) Raw264.7 were transfected with a luciferase reporter carrying WT or MUT-Nrp2 3′-UTR in the miR-328 mimics (H left) and miR-328 (H right) or their respective negative controls. *p < 0.05, **p < 0.01. Data are representative of at least three independent experiments.

To investigate osteoclastogenesis regulated by miR-328 in vitro. RAW264.7 cells were transfected with miR-328 mimics or inhibitors. The miR-328 expression levels were significantly upregulated by miR-328 mimic transfection and markedly inhibited by miR-328 inhibitor transfection (Figure 5B). Furthermore, overexpression of miR-328 suppress endogenous expression of OPG protein level (Figure 5C).

Consistent with the results of OPG protein expression, we found that miR-328 mimics enhanced TRAP staining, whereas miR-328 inhibitors weakened TRAP staining (Figure 5D). Overall, these data showed that osteoclast differentiation was promoted by miR-328 in vitro.

To determine bone resorption regulated by miR-328 in vivo. we injected miR-328 inhibitors or their negative controls into tail vein of 8-week-old male mice third per week for 8 weeks. Mice treated with miR-328 inhibitors showed increased trabecular bone volume, number, and thickness, and decreased trabecular separation relative to their negative groups. These data showed that bone resorption was regulated by miR-328 in vivo.

To investigate target mRNAs of miR-328 in osteoclast differentiation process, we have checked the miRNA-binding site in Targetscan and found that miR-328 binging site on Nrp2 3′-UTR (Figure 5G). Previously, Nrp2 deficiency has been reported to promote osteoclast differentiation and reduced osteoblast numbers^17^. Thus, luciferase reporter constructs containing the WT or mutated predicted miRNA-binding sites of Nrp2 were constructed. Then, RAW264.7 cells were transfected with WT-Nrp2–3′-UTR or MUT-Nrp2–3′-UTR with miR-328 mimics or negative controls and the luciferase activity of the Nrp2 reporter gene was inhibited by miR-328 mimics and the Nrp2 reporter gene was promoted by miR-328 inhibitors, whereas MUT-Nrp2–3′-UTR prevented this inhibition (Figure 5H). In conclusion, these data showed that miR-328 directly targets Nrp2 and inhibited Nrp2 expression in osteoclast cells.

### A549-Exos^miR-328 Inhibitors^ inhibited bone resorption *in vivo*

To investigate the therapeutic potential of miR-328 on bone metastasis, A549-Exos^miR-328 Inhibitors^ was generated. A549 cells transfected with miR-328 inhibitors derived exosomes were labeled with lipophilic dye Dio and assessed whether A549-Exos could carry miR-328 inhibitors and be transported to Raw264.7 cells. The fluorescence microscope revealed that A549-Exos^miR-328 Inhibitors^ could be successfully absorbed by Raw264.7 cells (Figure 6A). Then, we injected A549-Exos or A549-Exos^miR-328 Inhibitors^ into tail vein of 8-week-old male mice third per week for 12 weeks. Mice treated with A549-Exos^miR-328 Inhibitors^ showed increased trabecular bone volume, number, and thickness, and decreased trabecular separation relative to A549-Exos group, which showed that A549-Exos^miR-328 Inhibitors^ could alleviate bone loss caused by the bone metastasis (Figure 6B, 6C). In addition, TRAP staining showed fewer number of osteoclasts on the trabecular bone surface in A549-Exos^miR-328 Inhibitors^ groups than A549-Exos (Figure 6D, 6E), which revealed that an inhibitory role of A549-Exos^miR-328 Inhibitors^ in the differentiation of osteoclasts. In conclusion, these data suggested that intravenous injection of A549-Exos^miR-328 Inhibitors^ inhibited bone resorption in vivo and alleviate the osteopenia caused by bone metastasis.

**Figure 6.**
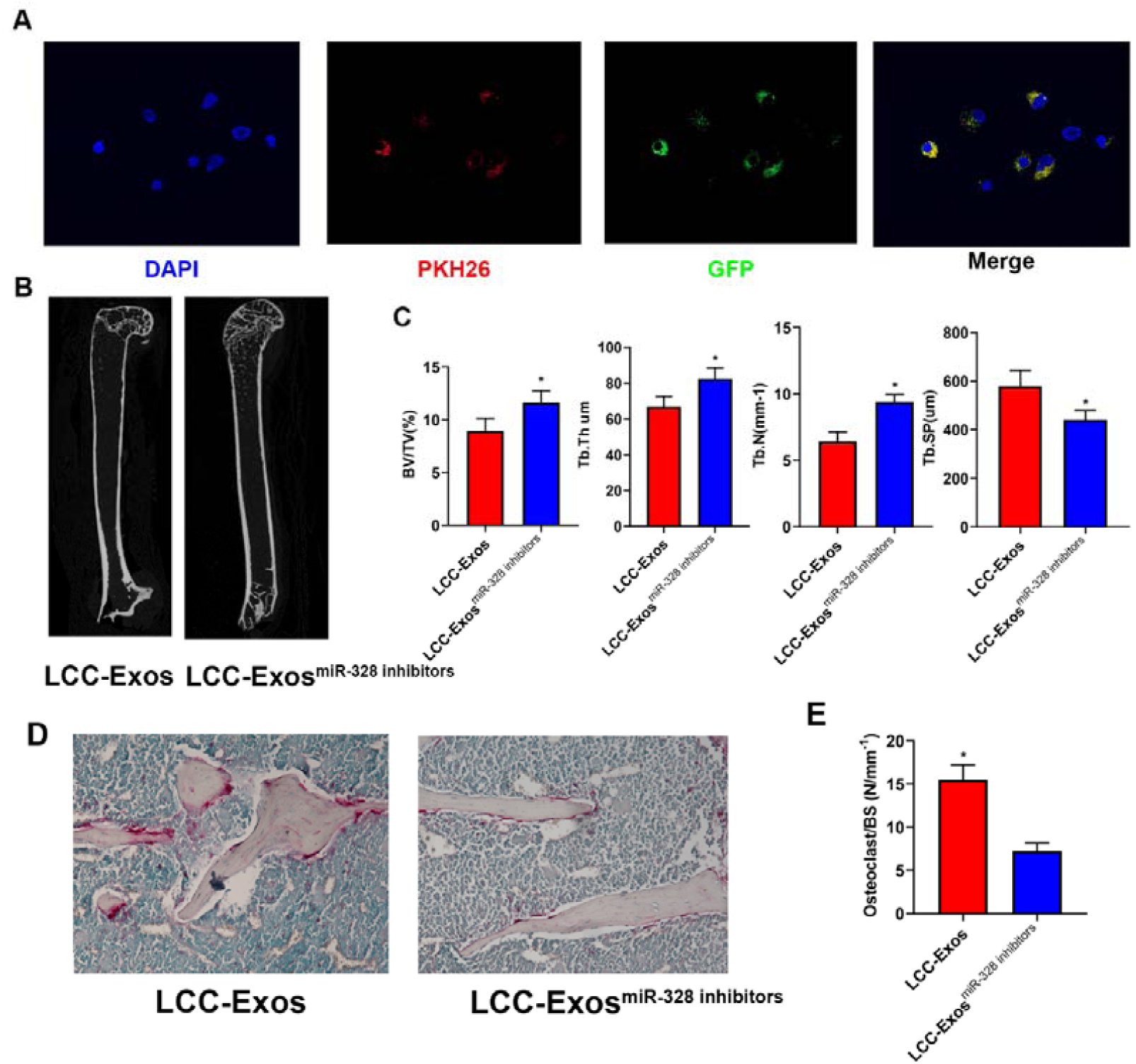
A549-Exos^miR-328 Inhibitors^ inhibited bone resorption *in vivo*. (A) Fluorescence microscopy analysis of RAW264.7 cells treated with DiO-labeled A549-Exos transfected with miR-328 inhibitors for 2h (B) Representative micro-CT images in femora from A549-Exos group and A549-Exos^miR-328 Inhibitors^ group (n = 6 per group). (C) Quantitative analysis of trabecular microarchitecture from A549-Exos group and A549-Exos^miR-328 Inhibitors^ group (n=8 per group). (D) Representative TRAP staining of femora from different groups (n=8 per group). (E) Quantitative analysis of osteoclasts per bone surface in different groups. (*P<0.05).

## Discussion

In this study, we found that osteoclast differentiation was enhanced when co-culturing with A549 cells. A549-Exos promote osteoclast differentiation in vitro. Furthermore, A549-Exos could be targeted for bone tissues and promoted bone resorption in vivo. Moreover, we found that A549-Exosomal miR-328 promoted bone resorption by targeting Nrp2. In addition, A549-Exos^miR-328 Inhibitors^ treatment in vivo ameliorated bone loss and prevented pathological fractures.

Previously, several cancer cells derived exosomes have been reported to be home to the bone microenvironment and cause metastatic lesions^18^. As murine prostate cancer line TRAMP-C1 inhibits the differentiation of the monocytic cell line RAW264.7 into osteoclasts^19^, non-small cell lung cancer (NSCLC) cells secrete exosomes promoted osteoclast differentiation of murine RAW264.7 cells by activation of EGFR phosphorylation^20^, exosomes released by multiple myeloma cells increase the viability and migration of osteoclast precursors, through the increasing of CXCR4 expression^21^ and extracellular vesicles isolated from PC3 culture medium are internalized into osteoclast precursors and osteoblasts^22^. In this study, A549 cell-derived exosomal miR-328 is the regulator of bone resorption and A549-Exos^miR-328 Inhibitors^ could ameliorate bone loss. Thus, A549 cells derived exosomes could load small molecule drugs and target bone tissues. Since a set of tumor cells derived exosomes could target bone tissues, we hypothesized that these exosomes can also be used as targeted drug carriers for the corresponding primary tumors. Our work might provoke interesting future works regarding the relationship between bone metastasis and tumor cells derived exosomes leading to new potential therapeutic targets.

A set of exosomal miRNAs has been reported to be correlated with bone formation or bone resorption^23^. For example, Exosomal miR-141-3p regulates osteoblast activity to promote the osteoblastic metastasis of prostate cancer^24^, overexpression of has□miR□940 in tumor exosomes and its ability to ultimately lead to extensive osteoblastic lesions in the resulting tumor^25^ and mi-192 cargo within exosome like vesicle transfer influences metastatic bone colonization^26^. These miRNAs may also serve as therapeutic potential targets for the treatment of bone metastases from lung cancers, which requires further investigation.

## Conclusions

Lung adenocarcinoma cell-derived exosomes (A549-Exos) promoted osteogenesis and bone resorption in vitro. Furthermore, A549-Exos target bone in vivo and promoted bone resorption in vivo. Mechanistically, A549-Exosomal miR-328 promote osteoclastogenesis by targeting Nrp2 and A549-Exos^miR-328 Inhibitors^ may serve as a potential nanomedicine for the treatment of bone metastasis.

## Acknowledgements

Not applicable.

## Author contributions

Conceived and designed the review: HBD and ZQW. Collected the data: ZYZ and CCZ. Analyzed the data: JY and WS. Supervised the study: JRQ and CCZ. Wrote the paper: CCZ, JRQ and LY.

## Availability of data and materials

Data sharing is not applicable to this article as no datasets were generated or analyzed during the current study.

## Ethics approval

All animal experiments were conducted in accordance with the National Institute of Health (NIH) Guide for the Care and Use of Laboratory Animals, with the approval of Longhua hospital (No.LHERAW-19038).

## Consent for publication

All authors reached an agreement to publish the study in this journal.

## Figure legends

**Figure S1.**
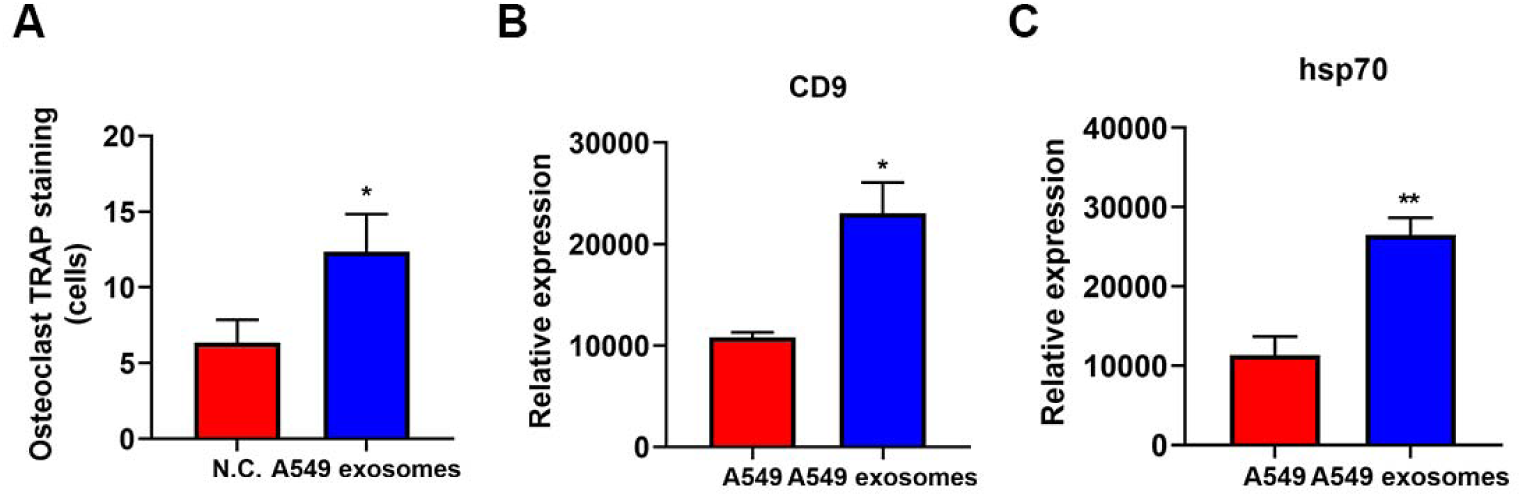
**(A)**. Quantitative analysis of positive TRAP staining osteoclasts in Figure 1A. (B). The quantification analysis of the Western Blot of CD9 and HSP70. *P < 0.05, **P < 0.01.

**Figure S2.**
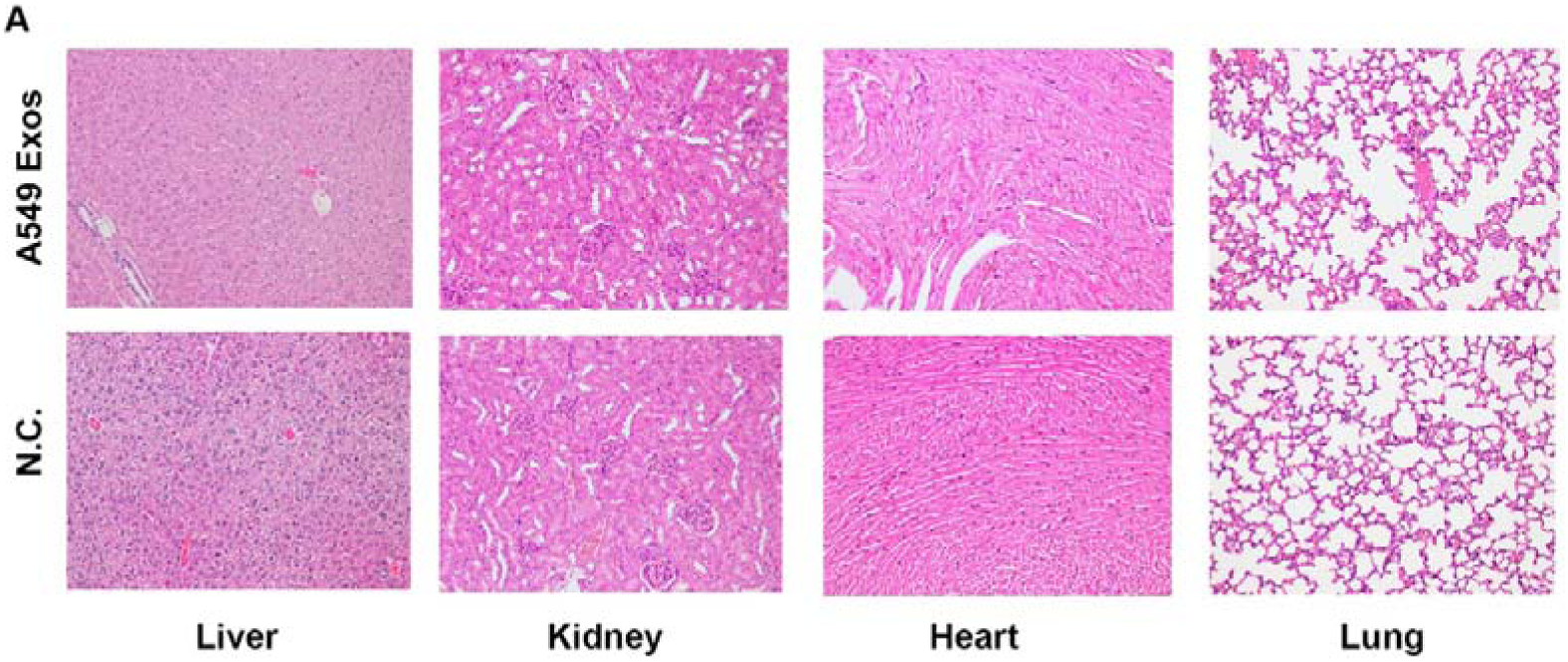
**(A)**. There was no histomorphometric change in the heart, liver, lung and kidney.

